# Speed-independent modulation of locomotor gait preference by sensory feedback in mice

**DOI:** 10.1101/2023.03.18.533262

**Authors:** Zane Mitrevica, Andrew J Murray

## Abstract

Locomotion is one of the most ubiquitous motor actions in the animal kingdom, essential for behaviours as diverse as foraging, migration, and escape. Successful execution of all these tasks relies on continual adjustment of locomotor gait in line with the behavioural demand for speed as well as the terrain. Failure in this process would disrupt locomotor smoothness, raise its energetic cost, and increase the risk of injury due to skeletal stress [1, 2]. Animals avoid these scenarios, in part, by transitioning from left-right alternating (walk, trot) to synchronous (gallop, bound) gaits as they increase the speed [3, 4]. However, this relationship is not deterministic [5, 6] and its connection to biomechanical factors, like the loading of limbs [7, 8], is unclear. To address this, we developed a head-fixed locomotor paradigm that decouples the speed- and leg loading-related influences on gait by combining optogenetic stimulation of an established speed-control pathway [9, 10] with head height or surface incline modulation. We found a pronounced speed-independent shift in homolateral limb coordination from strict alternation to a gallop-like pattern at upward oriented body postures and upsloping terrains. Both conditions are associated with greater relative loading of the hindlimbs and have a consistent effect on gait preference during head-fixed and head-free locomotion. These results suggest that mice use proprioceptive feedback from the limbs to coordinate their gait across speeds and environments, and implicate ipsilateral control mechanisms in this process. More broadly, our work serves as a principled entry point to a behaviour-driven study of gait circuits.

## Results

### Body tilt and surface incline modulate the anteroposterior distribution of leg loads

Selection of locomotor gait has been postulated to depend on the vertical ground reaction forces (GRF) animals experience while traversing paths of varying slopes and frictions [11, 12], carrying weights [7], or adjusting the tilt of the body [8]. To probe the influence of such sensory feedback on interlimb coordination in a principled manner, we sought to control the distribution of leg loads in an experimentally amenable species like the mouse. We hypothesised that this could be achieved in an ethologically relevant manner by modulating the body tilt of the animal or the surface incline. chose to explore both of these variables given their distinct internal and environmental origins respectively.

To characterise how changes in the body tilt and incline affect the loading of the limbs, we positioned head-fixed mice on four single-point load cells and measured the vertical GRFs exerted by individual legs during 5-second standstills (Figure 1A). To generate a wide range of body tilts with minimal intrusion, we manipulated the height of the animal’s head rather than its body tilt directly. Head height modulation over a 20-25 mm range at zero incline led to nonlinear changes in body tilt such that mice assumed more hunched postures at lower head heights and the relationship plateaued near the maximum head height compatible with a fully quadrupedal posture (*p*_*linear*_ = 2.9 × 10^−60^, *p*_*quadratic*_ = 1.2 × 10^−5^ *t-test*; data from 7 mice; Figure 1B). It also resulted in a systematic redistribution of body weight between the limbs and the head-fixation apparatus (Figures S1A,B). Specifically, tilting the body upward reduced the weight borne by the forelimbs in a linear manner, by 2.5 ± 0.1% of body weight per degree of body angle (*p*_*f ore*_ = 9.3 × 10^−42^; Figure 1C). This was due to a transfer of weight onto the head-fixation system, with no significant effect on the hindlimb load (*p*_*head*_ = 6.2 × 10^−41^, *p*_*hind*_ = 0.4 *t-test*). In contrast, changing the surface incline over an 80 degree range, from a downward to an upward slope, increased the absolute loading of the hindlimbs irrespective of the head height (*p*_*hind−linear*_ = 2.6 × 10^−73^, *p*_*hind−quadratic*_ = 1.8 × 10^−9^ *t-test*; data from 7 mice; Figure 1D). This occurred primarily at the expense of the forelimb load, while interactions with the head-fixation apparatus, although non-negligible, were highly variable across mice (*p*_*f ore−linear*_ = 9.5 × 10^−36^, *p* _*f ore−quadratic*_ = 6.0 × 10^−18^ *t-test*). These data therefore highlight fundamental differences in how the body tilt and surface incline affect the absolute limb GRFs at least under head-fixed circumstances.

**Figure 1:**
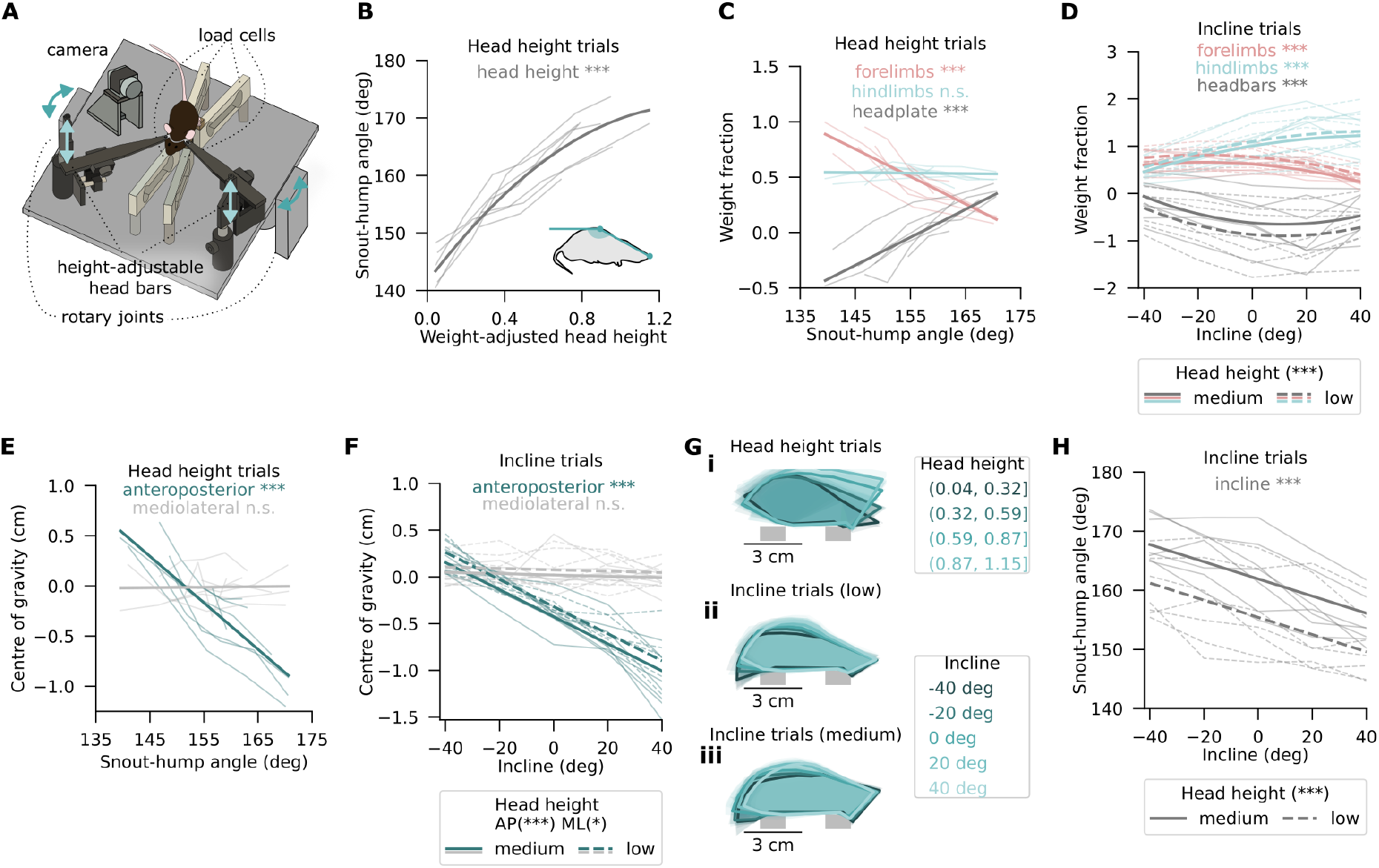
Body tilt and surface incline modulate the anteroposterior distribution of leg loads. **(A)** Schematic representation of the GRF measurement setup that allows modulation of mouse head height (light arrows) and incline (dark arrows). **(B)** Body tilt quantified as the obtuse angle between the snout-hump vector and the ground (inset) and plotted as a function of weight-adjusted head height of individual mice across trials on zero incline (thin lines) and as a second order polynomial mixed-effects model fit across mice (thick, model statistics in Table S1). **(C)** Fraction of body weight placed on the forelimbs (pink), hindlimbs (blue), and the head-fixation apparatus (grey) during the same standstill trials as in (B). Trial averages of individual mice computed over five discrete body tilt ranges are displayed by the thin lines, while the thick lines are linear mixed-effects model fits. Negative values reflect a “weight gain” due to the normal force at the head/head-bar interface. **(D)** Same as (C) but over a range of inclines at a higher (solid lines) and lower (dashed) head height 7 mm apart. **(E)** Same as (C) but showing the centre of gravity across limbs along the mediolateral (grey) and anteroposterior (teal) body axes. Zero corresponds to equal weight placed on forelimbs and hindlimbs. **(F)** Same as (E) but over a range of inclines as in (D). **(G)** Body posture of individual mice (shaded polygons) and across mice (thick outlines) at four ranges of weight-adjusted head heights (*i*) and five inclines (*ii, iii*). The shapes are aligned at the back corner of the front support post (approximated in grey). **(H)** Same as (B) but over a range of inclines as in (D).

Previous work on GRF involvement in gait control has not explicitly linked the absolute loading of the legs to interlimb coordination, leaving open the possibility that the relative load on the limbs or other biomechanical factors have a causal relation to gait [13]. In our paradigm, upward slopes and body orientations both increased the hindlimb load relative to that of the forelimbs, shifting the centre of gravity anteroposteriorly by 1.5 ± 0.1 mm*/*deg*ground* and 4.6 ± 0.2 mm*/*deg*body* respectively (*p*_*AP−ground*_ = 1.7 × 10^−46^, *p*_*AP−body*_ = 4.9 × 10^−52^ *t-test*; Figures 1E, 1F, S1C, S1D). Over the entire range of examined inclines and body tilts, we recorded 0.6-1.6 cm anteroposterior shifts in the centre of gravity, with no significant difference between trial types and no change along the mediolateral body axis (*p*_*AP−trialtype*_ = 0.14 *one-way ANOVA*; *p*_*ML−ground*_ = 0.05, *p*_*ML−body*_ = 0.8 *t-test*). Notably, the incline-dependent redistribution of weight was actively countered by mice through changes in body tilt and foot placement (Figure 1G). Like several other species moving on inclined surfaces [14–17], head-fixed mice adopted more hunched postures and positioned their feet more anteriorly when standing on upward slopes (*p*_*body−tilt*_ = 2.2 × 10^−5^, *p* _*f oot−position*_ = 0.006 *t-test*; Figures 1H, S1E). Altogether, these results suggest that self-driven adjustments of body tilt during head-fixed locomotion, like external changes in the surface slope, systematically modulate the relative vertical GRFs of the limbs as well as the body’s centre of gravity (Figure S1H).

### Changes in body tilt and surface incline shift the homolateral limb phase

Animals as diverse as horses and cockroaches have been found to switch from left-right alternating to out-of-phase stepping patterns at lower speeds when bearing weights or moving on a slippery surface [7, 11]. These two conditions have in common an increased impact of the vertical GRF component. However, the critical behavioural variable could still be the absolute loading of the limbs, the GRF distribution across the limbs, or even posture on its own. We aimed to tease these possibilities apart by combining incline and head height manipulations, given their differential effects on biomechanical and postural variables. Specifically, we head-fixed mice atop a passive treadmill and modulated their head height or treadmill incline, similar to the previous experiment on the force sensors (Figure 2A). To elicit locomotion of wide-ranging speeds and gaits with consistency, we expressed channelrhodopsin in the glutamatergic neurons of the cuneiform nucleus (CnF), a region known to trigger locomotion at a speed that scales with stimulation intensity [9, 10, 18, 19] (Figure 2B, 2C, S2A).

**Figure 2:**
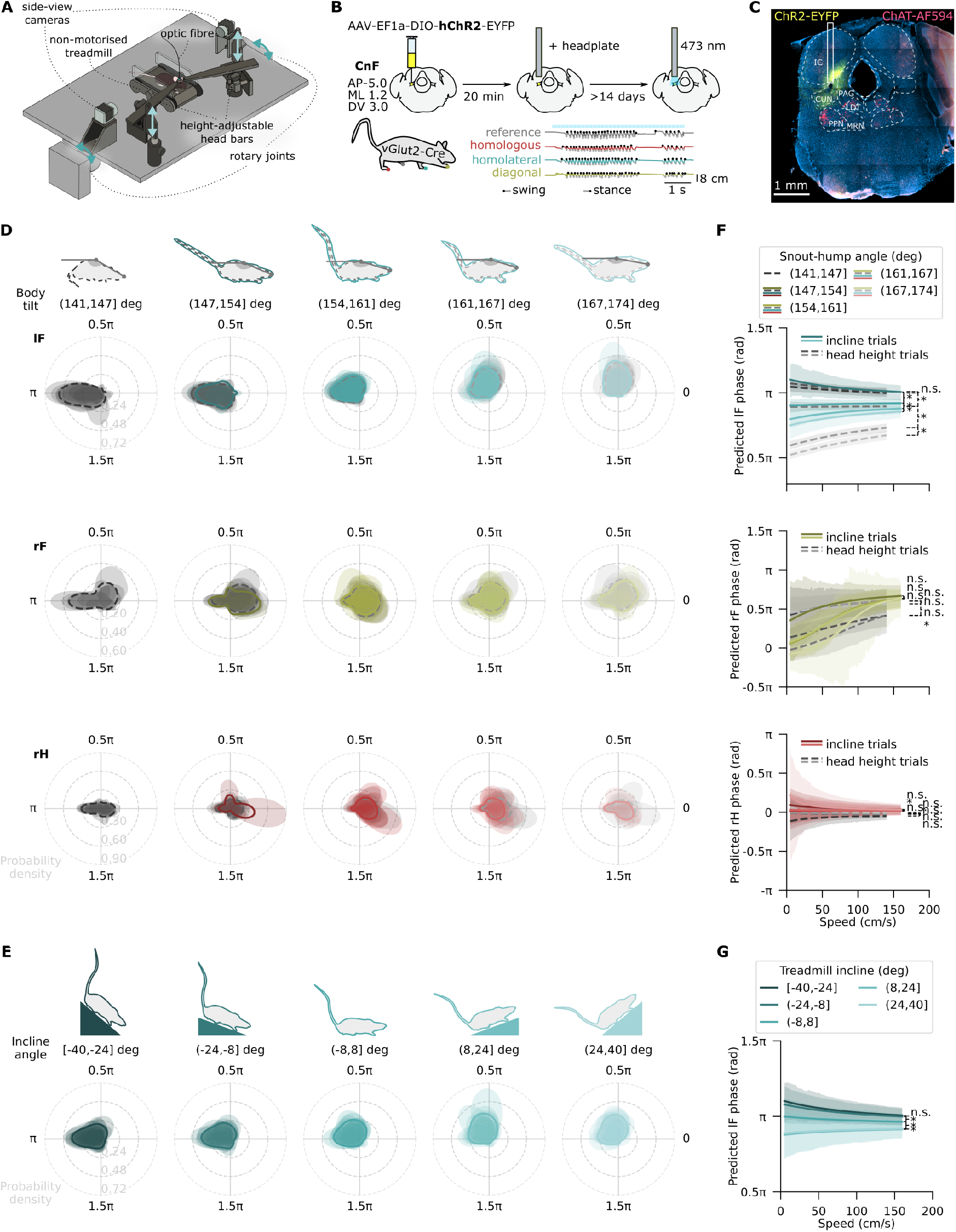
Changes in body tilt and surface incline shift the homolateral limb phase. **(A)** Illustration of the passive treadmill setup that allows modulation of mouse head height (light arrows) and incline (dark arrows). **(B)** *Top and bottom left*: experimental strategy and timeline. *Bottom right*: example x-coordinate traces of the four limbs tracked by DeepLabCut during a 50 Hz optogenetic stimulus (grey shaded region; light pulses shown by the blue bars above). Black and dark grey dots mark the detected swing and stance onsets respectively. **(C)** An example coronal section showing the site of virus injection in yellow, choline acetyltransferase (ChAT) antibody staining in red, and optic fibre implantation. **(D)** Density distributions of left forelimb (lF, top), right forelimb (rF, middle), and right hindlimb (rH, bottom) phases relative to the reference limb (left hindlimb, lH) across five intervals of snout-hump angles (semi-transparent distributions), with the same mice (n=12) completing both incline (teal/yellow/red, solid lines) and head height (grey, dashed lines) trials. Average distributions across mice are shown by solid lines. Limb phase values of 0 and *π* reflect synchrony and alternation respectively. Data from mice with fewer than 40 strides per category were excluded from the respective plot. **(E)** Same as (D) but across five intervals of treadmill inclines and for lF only. **(F)** lF (top), rF (middle), and rH (bottom) phases relative to lH as a function of speed grouped by snout-hump angle as in (D) and shown as fits of a Bayesian mixed-effects model for circular data (solid lines) with 95% highest posterior density intervals (shaded regions). **(G)** Same as (F) but grouped by incline and for lF only. Statistical significance in Bayesian mixed-effects models (F,G): * significant, n.s. not significant.

CnF stimulation at 10-50 Hz reliably induced locomotion at short latencies (*<* 150 ms in 88% trials), with the running speed linearly related to stimulation frequency (*p*_*median−speed*_ = 7.7 × 10^−42^ *t-test*; *p*_*max−speed*_ = 3.3 × 10^−34^; Figure S2B). Importantly, optogenetically induced locomotion comprised a wide range of limb coordination patterns. The average distributions of limb phases were similar to idealised distributions of trot and bound, likely reflecting a mix of alternating, synchronous, and transitional gaits (Figures S2C, S2D). This was further evidenced by a variety of limb support motifs displayed at all speeds, and was in contrast to non-stimulated locomotion on a motorised treadmill, which was dominated by trot (Figures S2D, S2E). Lastly, kinematic parameters, like stride length and frequency, were not significantly different during stimulated head-fixed and self-initiated unrestrained locomotion (Figures S2F, S2G). We therefore considered this locomotor paradigm suitable for studying naturalistic locomotion across the full spectrum of mouse gait while controlling the GRF distribution.

Over a period of two weeks, mice (n=12) performed both head height and incline trials as we explored the impact of body tilt and surface slope on limb coordination. Changes in body tilt over a 33 degree range revealed a pronounced shift in phase preference of the left forelimb (lF) relative to the left hindlimb (lH). At hunched postures (intervals (141,147], (147,154] deg), these homolateral limbs moved predominantly in strict alternation, while more upward-oriented body tilts (intervals (161,167], (167,174] deg) showed a prevalence of out-of-phase coordination (permutation test, *p <* 0.0001; grey in Figures 2D, statistics in S2H). Such a notable transition in the relative phase preference was not seen for either the right forelimb (rF) or hindlimb (rH), but was mirrored by treadmill incline modulation (Figures 2D, 2E, S2H). In the incline trials, the fixed absolute head height narrowed the range of adopted body tilts, yet downward body and treadmill orientations both favoured strict homolateral alternation, while out-of-phase coordination dominated in the two most upward oriented body tilt and incline intervals (permutation tests *p <* 0.0001; teal in Figures 2D, 2F; statistics in S2H, S2I, S2J). Viewed in isolation of other limbs, these lF phase shifts resembled a transition from trot-like homolateral coordination to one characteristic of right-leading transverse gallop (Figures S2C, S3A). However, gait selection is also known to be influenced by locomotor speed and the descending pathway from the CnF that at least partly mediates it [5, 9, 10]. To explicitly account for this influence, we fit a circular mixed-effects model [20] to the limb phase data with speed and snout-hump angle intervals as fixed predictors. Speed had a significant impact on the lF phase as expected, yet the impact of body tilt was independent of speed (Figures 2F, 2G Table S1). Analogous models of the two other limb pairs also revealed a significant influence of speed but not the snout-hump angle in line with the previous analyses (Figure 2F, Table S1). Altogether, these data suggest a similar influence of speed on all limb pairs, while pointing to homolateral coordination as the primary target of body tilt and incline modulation.

### Head-free locomotion supports a sensory feedback-dependent shift in homolateral coordination

So far, we have shown a pronounced shift in homolateral coordination upon changes in body tilt and treadmill incline during optogenetically evoked, head-fixed locomotion. To determine whether these relationships persist during self-initiated, head-free movement, we recorded mice on a levelled or sloped motorised treadmill (n=12 and n=10 mice respectively, two separate cohorts; Figure 3A). Despite the dominance of left-right alternating gaits under these circumstances (Figure S2C, S2E), the range of observed homolateral phases was broad enough to enable pairwise comparison of body tilt angle distributions across limb phase intervals spanning [0, 1.6*π*]. We found a markedly greater prevalence of upward oriented body postures during out-of-phase homolateral movement (interval (0.4*π*, 0.8*π*]) than during alternation (interval (0.8*π*, 1.2*π*]), which precisely mirrors the relationship seen on the passive treadmill (Figure 3B, 3C, S3B). At lower speeds, upward body tilts were also common when an lF step followed shortly after lH as in lateral walk (interval [0, 0.4*π*]). Both phase-body tilt associations were independent of speed (interaction HPD = (−2.3,2.3); Figure S3C) and persisted when averaged across various inclines (teal in Figures 3B, 3C; Figure S3C). Furthermore, the upward slopes were also associated with progressively out-of-phase homolateral coordination akin to head-fixed locomotion (Figures 3D, S3D). The effects of both body tilt and incline on the homolateral stepping pattern were, on average, smaller than on the passive treadmill (Figures 3E, 3F). While this reflects the reduced propensity to switch gait in the absence of CnF stimulation, the inability to modulate leg load through interaction with the head-fixation apparatus and the greater postural adjustments made to oppose changes in the surface slope might also contribute (Figure S3E, S3F). These results are consistent with limb GRFs having a role in gait selection and speak to the ethological relevance of body tilt and incline in sensory-dependent control of locomotion.

**Figure 3:**
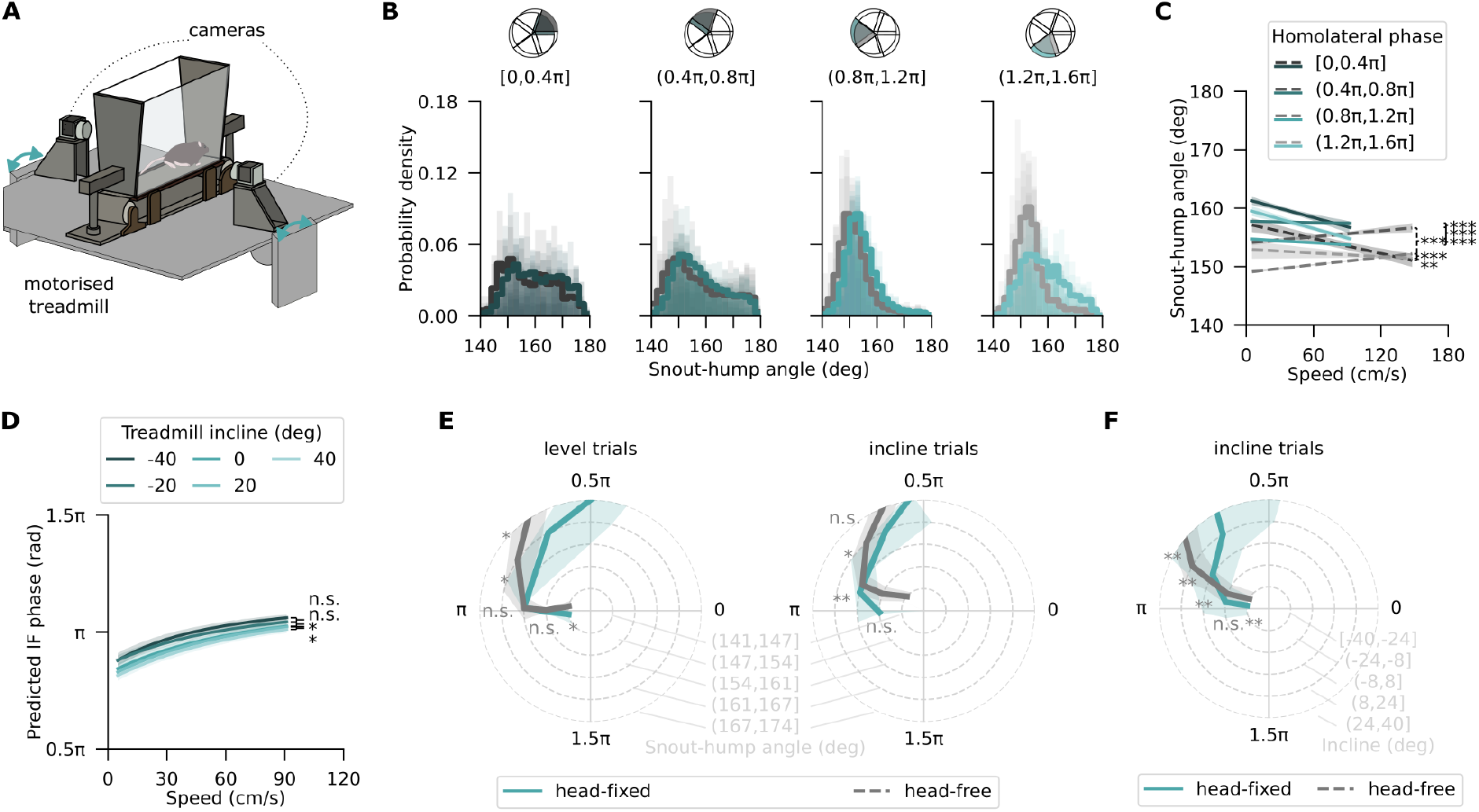
Head-free locomotion supports a sensory feedback-dependent shift in homolateral coordination. **(A)** Illustration of the motorised treadmill setup that allows incline modulation (dark arrows). **(B)** Density distributions of snout-hump angles across four categories of relative lF phases (semitransparent distributions), with the average distributions across mice shown by solid lines. Limb phase values of 0 and *π* reflect synchrony and alternation respectively. Data from mice with fewer than 40 strides per category were excluded from the respective plot. Only one mouse met this criterion in phase interval (1.6*π*, 2*π*], hence it was not plotted. **(C)** Snout-hump angles as a function of speed grouped by lF phase as in (B) and shown as fits of a linear mixed-effects model (solid lines) with 95% confidence intervals (shaded regions). **(D)** lF phases relative to lH as a function of speed grouped by treadmill incline, shown as fits of a Bayesian mixed-effects model for circular data (solid lines) with 95% highest posterior density intervals (shaded regions) **(E)** Peak lF phases relative to lH grouped by snout-hump angle during head-fixed, optogenetically stimulated (teal) and head-free, non-stimulated (grey) locomotion on levelled (*left*) and sloped(*right*) surface. Shaded regions show standard deviation. **(F)** Same as (E) but grouped by surface incline. Statistical significance in Bayesian mixed-effects models (D): * significant, n.s. not significant. Statistical significance thresholds in other panels prior to Bonferroni correction: * *p <* 0.05, ** *p <* 0.01, *** *p <* 0.001.

### Incline mimics the influence of body tilt and may be linearly additive

The sensory signal linking body tilt and incline to locomotor gait could, in principle, be vestibular, neck proprioceptive, GRF-related limb proprioceptive, or some combination of these. We consider direct vestibular involvement improbable because neither the linear acceleration due to gravity nor the tilt of the head was affected by our head height manipulation [21]. Conversely, proprioceptive signals from the limbs and neck were both likely modulated by changes in the body tilt, leg loading, or both. To separate these variables, we fit a series of Bayesian mixed-effects models with combinations of speed, body tilt, incline, weight, sex, and the choice of reference limb as predictors (Table S1). This revealed three arguments in favour of GRF redistribution being a relevant factor in gait specification. First, greater body mass was found to favour homolateral alternation just as heavier mice would be expected to require a greater redistribution of weight to attain a given leg-loading difference along the anteroposterior body axis (Figures S3G, S3H). Second, equal changes in body tilt corresponded to a greater homolateral phase shift during head height manipulation than in the incline trials, suggesting that body tilt and incline might influence gait through a common mechanism (Figure 4A). Finally, body tilt interacted with treadmill incline (*HPD*_*SSDO*_ = (−0.81, −0.63)) such that their respective influences on homolateral coordination were smaller at upward oriented treadmill and body configurations (Figure 4A). A significant interaction was also found between body tilt and speed, albeit only in the head height trials (*HPD*_*SSDO*_ = (0.60, 0.77); Figure 4A). This likely reflected a shared correlation with the head height itself rather than a biological phenomenon, and was therefore not considered further (Figures S3I, S3J). Altogether, our modelling suggests that incline and body tilt influence gait through a common mechanism and point to the leg load transducing system as the most relevant candidate.

**Figure 4:**
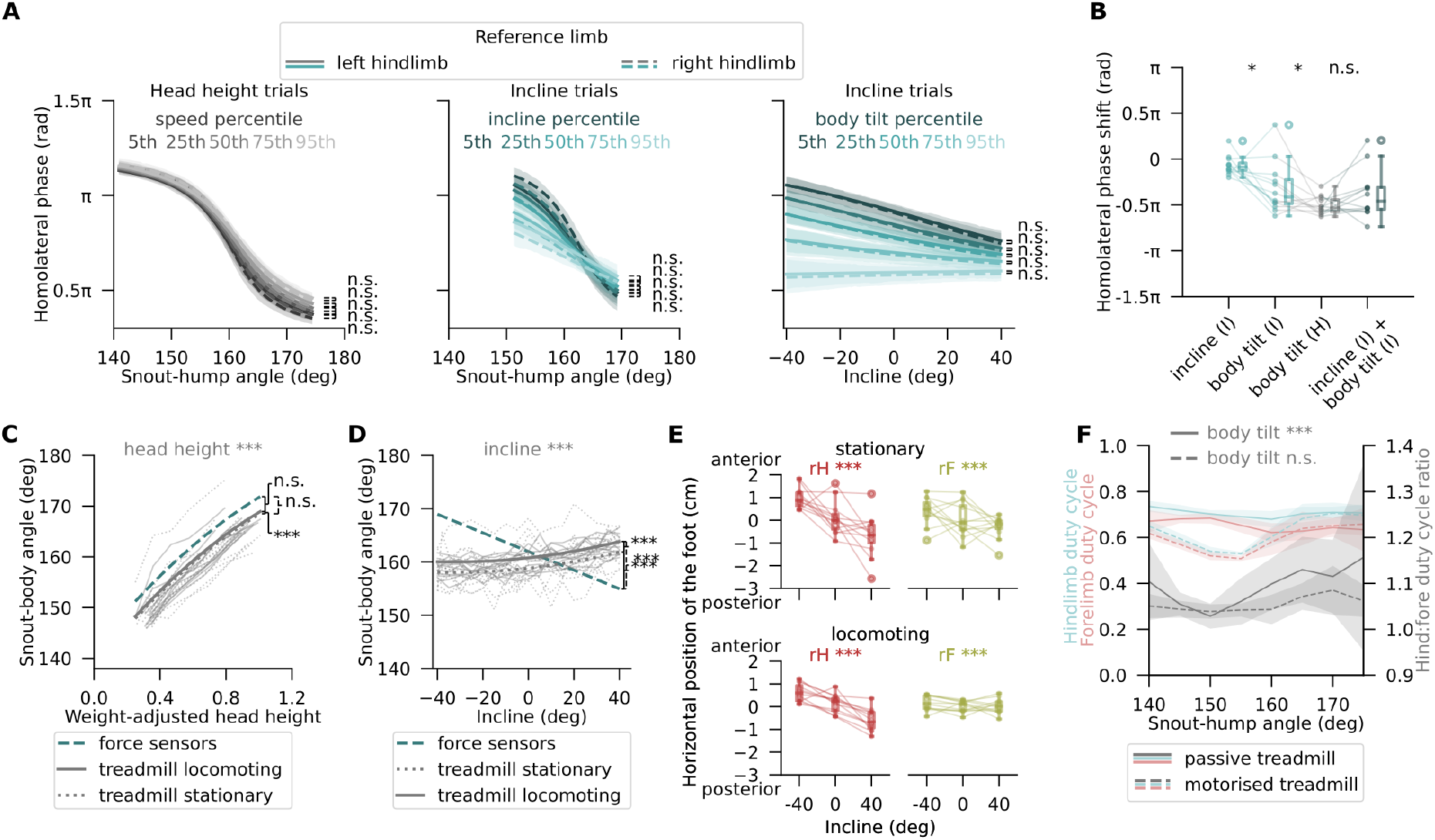
Incline mimics the influence of body tilt and may be linearly additive. **(A)** Forelimb phases relative to the homolateral hindlimb as a function of treadmill incline or snout-hump angle with lH (solid lines) and rH (dashed lines) as the reference limb. Shown are Bayesian mixed-effects model fits to the head height (*left*) and incline (*middle, right*) trial data at five percentiles of speed (*left*), inclines (*middle*), or body tilts (*right*) – parameters that significantly influence the effect of the respective independent variable. Shaded regions are 95% highest posterior density intervals. **(B)** GRF-equivalent homolateral phase shift due to body tilt and treadmill incline in incline trials (I; light teal) and head height trials (H; grey), and the combined GRF-equivalent phase shift in the incline (I; dark teal). **(C)** Body tilt angle as a function of weight-adjusted head height during optogenetically induced locomotion (solid) and the 500 ms pre-stimulus period (dotted) on zero incline, plotted for individual mice across trials (thin lines) and as linear mixed-effects model fits across mice (thick). The dashed curve shows a model fit to analogous data on the force sensors. **(D)** Same as (F) but over a range of inclines at a fixed medium head height. Linear coefficients: force sensor *β* = -0.15± 0.01 *deg*_*bodytilt*_*/deg*_*incline*_, treadmill *β* = 0.046 ± 0.002 *deg*_*bodytilt*_*/deg*_*incline*_. **(E)** Horizontal foot position depending on the treadmill incline during the 500 ms pre-stimulus period (*top*) and optogenetically induced locomotion (*bottom*). Linear coefficients: force sensor *β* = (1.206± 0.005) × 10^−2^ *cm/deg*_*incline*_, treadmill *β* = -(1.958± 0.008) × 10^−2^ *cm/deg*_*incline*_. **(F)** Average hindlimb (teal) and forelimb (pink) duty cycles, and their ratio (grey) during head-fixed (*p*_*ratio*_ = 5.5 × 10^−16^, one sample t-test) and head-free (*p*_*ratio*_ = 0.58) locomotion, across snout-hump angles. Mean values across mice are shown, along with 95% confidence intervals. Statistical significance in Bayesian mixed-effects models (A): * significant, n.s. not significant. Statistical significance thresholds in other panels prior to Bonferroni correction: * *p <* 0.05, ** *p <* 0.01, *** *p <* 0.001.

To probe this further, we used force sensor data (Figures 1D, 1F) to compute GRF-equivalent phase shifts attributable to changes in body tilt or treadmill incline on a per-mouse basis (see Methods). If GRF was the sole important variable, we would expect the effects of incline and body tilt to be linearly additive up to a point of saturation. Equated in the GRF space, the sum of incline- and body tilt-related phase shifts during the incline trials was indeed indistinguishable from the phase change seen in the head height trials (Figure 4B). This result is consistent with body tilt having no effect on homolateral coordination beyond that mediated through GRF redistribution. However, it relies on incline and body tilt having setup-invariant effects on the distribution of leg loads, which could not be directly verified. Using postural variables as a surrogate measure, we found no setup-dependence of the relationship between body tilt and the height of the head, implying that the indifference assumption may hold in the head height trials (*p*_*standing*_ = 0.05; *p*_*locomoting*_ = 0.09; Figure 4C). At the same time, the strong correlation between body tilt and incline seen on the force sensors was greatly reduced on the treadmill, and incline modulation was primarily opposed through foot displacement instead (Figures 4D, 4E). Given this change in the adaptive strategy, the apparent additive effects of incline and body tilt should be treated with caution, and a contribution of neck proprioceptors to gait selection, in addition to a GRF-dependent mechanism, cannot be excluded.

Beyond sensory processing, the feedback-dependence of gait could either be implemented by interlimb neural coupling or emerge from intralimb changes in the stance-to-swing timing. In the latter case, an anteroposterior shift in the centre of gravity might be expected to alter the timing of interlimb coordination by selectively prolonging the stance phase of more heavily loaded legs [22]. Indeed, at upward body and treadmill orientations during head-fixed locomotion, we found an increase in the hindlimb duty cycle, namely the stride fraction spent in stance, relative to that of the forelimbs (Figures 4F, S3K). This result was most pronounced during head-free locomotion on an inclined surface such that every 10 degree shift in the treadmill slope, on average, changed the hindlimb-to-forelimb duty cycle ratio by 4% (p = 5.9 × 10^−15^ *t-test*; Figure S3K). At the same time, the effects observed upon incline modulation under head-fixed conditions and freely moving changes in body tilt were an order of magnitude smaller. Such inconsistent results across the studied locomotor settings suggest that the observed modulation of gait by sensory feedback is unlikely to emerge solely from intralimb mechanisms even though they might play a role.

## Discussion

We have shown that the choice of locomotor gait in mice, specifically the coordination of their homolateral limbs, strongly relates to the anteroposterior position of the animals’ centre of gravity. Whether the latter is displaced by factors of internal (body tilt) or environmental (surface incline) origin, an increased loading of the hindlimbs invariably coincides with a speed-independent shift in the homolateral phase preference from strict alternation to out-of-phase coordination. This effect resembles a trot-to-gallop transition, is robust across a range of locomotor conditions, and supports a gait control mechanism that relies on a moment-by-moment comparison of the vertical GRFs borne by the legs.

A GRF-based approach to gait selection has been previously proposed as a temporally feasible alternative to a strategy rooted in energy cost minimisation [7] and has featured in theoretical work on quadrupedal locomotion [8, 23–25]. While such models are yet to cohesively capture the transition from trot to gallop, empirical studies of load-bearing dogs and horses have linked rearward shifts in the body weight distribution during trot to a greater galloping propensity [7, 26, 27]. Moreover, quadrupeds with naturally sloped backs, like giraffes, hyenas, and gnus, avoid trot altogether in favour of gallop-like gaits [28]. Our study provides direct experimental support for these ideas by revealing similar effects on interlimb coordination when GRFs are manipulated through body tilt and surface incline. It also provides the first systematic demonstration of feedback-dependent gait control in a non-cursorial quadruped. This is a nontrivial finding given that the skeletal structure and movement kinematics of mice are fundamentally different from those of cursorial species [29, 30] and could have led to alternative gait control principles.

The widespread prevalence of this particular gait selection strategy across animal phyla suggests that it might provide unique advantages. One hypothesis is that GRFs borne by the limbs are directly proportional to bone stress, and the switch from trot to gallop reduces both of these variables [1, 7]. A change in gait can therefore serve to prevent injury and ensure skeletal integrity at higher locomotor speeds than allowed by trot. In fact, animals differing in size by over four orders of magnitude experience similar peak bone stresses at equivalent speeds and gaits [2, 31], which highlights the resilience of GRF-based gait control against evolutionary pressures. It does not, however, explain why it is the anteroposterior distribution of GRFs, not their absolute magnitude, that seems to be of importance. One appealing hypothesis attributes this phenomenon to load-dependent regulation of limb duty cycles that, when substantially asymmetrical, cannot mechanically support the diagonal limb synchrony characteristic of trot [8]. While aspects of this idea have been supported by empirical work [22, 26, 27], the effects of GRFs on stance duration we observed were inconsistent across locomotor conditions and unlikely to account for the feedback-dependent shift in gait preference in its entirety. An alternative, albeit not mutually exclusive, perspective on gait selection focuses on the body pitch moment of inertia that an anteroposterior skew in the GRF distribution may induce and that is similar to the body tilt we have studied here. This hypothesis proposes that quadrupeds switch from trot to a three- or four-beat gait when the energy cost of vertically translating their centre of mass exceeds the cost of pitching [23, 24, 32]. Although energy optimisation does not seem to be the primary objective of gait transitions [7], it reflects a potential advantage of having the body tilt and gait to be closely coupled as implied by our results. Animals capitalise on this relationship during locomotion by flexibly adjusting their body tilt to attain target speeds or counteract the GRF redistribution imposed by the terrain [17, 33]. Our results cannot exclude the possibility of posture having an additional effect on gait independent of that mediated by the GRFs. It might also explain previous reports of the opposite, if any, relationship between interlimb coordination and surface incline, despite the leg load distribution being governed by the latter [15, 34].

A major contribution of our work is establishing a locomotor paradigm that reliably exposes the role of sensory feedback in gait selection, using an experimentally tractable species. While the choice of gait is well-known to be regulated by the descending speed-related signal from the CnF [9, 10, 18, 19], our research paves the way for a principled investigation of the neural circuits behind its GRF-dependence and the implementation of different gaits as such. For example, we have found that changes in the anteroposterior GRF distribution primarily affect the coordination of the homolateral, but not diagonal or homologous, limb pairs. This suggests that the speed-agnostic GRF-dependence of gait we report might be mediated, in part, by the propriospinal neurons which connect the ipsilateral limb segments in the spinal cord and have been previously implicated in speed- and context-dependent phase modulation of the homologous limbs [12, 35]. Future circuit-level work is required to reconcile these results with our behavioural findings, as well as to probe the causal involvement of proprioceptive afferents in feedback-dependent gait selection. Another exciting target of future inquiry might be a recently characterised neural population in the dorsal spinal cord that receives joint proprioceptive and cutaneous input [36]. Our paradigm opens the way to the kind of empirical work required to unravel the neural circuit basis of gait control.

## Acknowledgments

This work was funded by a Sainsbury Wellcome Centre Core grant from the Gatsby Charitable Foundation (219627/Z/19/Z to A.J.M.) and a Sainsbury Wellcome Centre PhD studentship (to Z.M.). We thank F.Claudi, F.Marbach, A.Akrami, E.Chong, R.Brownstone, M.R.Carey, M.W.Mathis, M.Stephenson-Jones, and members of the Murray laboratory for discussions, training, and support, and are grateful to T.Branco, M.Stephenson-Jones, and R.Brownstone for comments on the manuscript.

## Author contributions

Z.M. designed the study, performed all experiments and analyses, and wrote the manuscript. A.J.M. supervised the work.

## Declaration of interests

A.J.M. is a co-founder and the chief executive officer of Sania Therapeutics.

## Experimental procedures

### Resource availability

| REAGENT OR RESOURCE | SOURCE | IDENTIFIER |
| --- | --- | --- |
| **Antibodies** |  |  |
| Goat anti-ChAT | Millipore | Cat# AB144P, RRID: AB_2079751 |
| Alexa Fluor® 594 AffiniPure Fab Fragment Donkey Anti-Goat IgG (H+L) | Jackson Immunoresearch | Cat# 705-587-003, RRID: AB_2340435 |
| **Bacterial and virus strains** |  |  |
| AAV1-EF1a-DIO-hChR2(H134R)-EYFP | Addgene | Cat# 20289; RRID: Addgene_20298 |
| AAV2-EF1a-DIO-iChloc-2A-dsRed | Addgene | Cat# 70762; RRID: Addgene_70762 |
| **Experimental models: Organisms/strains** |  |  |
| Mouse: C57BL/6J | Charles River |  |
| Mouse: Vglut2-ires-Cre | Jackson Laboratory | JAX: 016963, RRID: IMSR_JAX:016963 |
| **Software and algorithms** |  |  |
| Analysis Python and R code | GitHub | https://github.com/zanemit/sensory-dependent-gait |
| Bonsai | Lopes et al. 2015 [37] | https://bonsai-rx.org |
| DeepLabCut | Mathis et al. 2018, Nath et al. 2019 [38, 39] | https://github.com/DeepLabCut/DeepLabCut |
| **Other** |  |  |
| Low-Friction Rodent-Driven Belt Treadmill | Jackson et al. 2018 [40] | Janelia 2017-049 |

#### Lead contact

Further information and requests for resources and reagents should be directed to and will be fulfilled by the lead contact, Zane Mitrevica.

#### Materials availability

This study did not generate new unique reagents.

#### Data and code availability

All data reported in this paper will be shared by the lead contact upon request.

Analysis code can be found at the following GitHub repository: https://github.com/zanemit/sensory-dependent-gait.

Any additional information required to reanalyse the data reported in this paper is available from the lead contact upon request.

### Experimental model and subject details

#### Animals

All experiments were performed under the Animals (Scientific Procedures) Act of 1986 (PPL PE4FA53CB) following local ethical approval. Male and female adult mice were housed on a reversed 12 hour light-dark cycle with ad-libitum access to food pellets and water. The passive treadmill and force sensor experiments were done with 10-20 weeks old Vglut2-ires-Cre mice (Jackson Laboratory, stock 016963). The motorised treadmill experiments used 6-9 weeks old Vglut2-ires-Cre and C57BL/6J wild type (Charles River 027) mice. Animals were weighed at least once a week during all behavioural procedures.

### Method details

#### Viruses

For optogenetic triggering of locomotion, 23 nL of AAV1-EF1a-DIO-hChR2(H134R)-EYFP (titer 1.0 × 10^13^, Sainsbury Wellcome Centre Viral Vector Core, henceforth SWC VVC, from Addgene plasmid 20289) was injected unilaterally into the cuneiform nucleus (CnF) of Vglut2-Cre mice at AP -5.0, ML 1.2, DV -3.0 mm from bregma. Control mice received the same injection but were of the C57BL/6J phenotype. These mice had no perceptible reaction to the optical stimulus and were not used for data acquisition.

A subset of Vglut2-Cre and wild type mice used in the motorised treadmill experiments had received a bilateral CnF injection of 23 nL AAV2-EF1a-DIO-iChloc-2A-dsRed (titer 1.1 × 10^14^, SWC VVC from Addgene plasmid 70762), but showed no discernible reaction to the optical stimulus, did not differ in their locomotor performance (one-way ANOVA, *p* = 0.29), and were therefore pooled for analysis.

#### Surgical procedures

Mice were anaesthetised with 4% isoflurane and maintained under anaesthesia for the duration of the surgery with 1.5-2.5% isoflurane (both in oxygen 1 L/min). Analgesia was provided through subcutaneous administration of meloxicam (5 mg/kg).

Viral vectors were delivered via pulled 3.5” glass pipettes (Drummond scientific) at a speed of 23 nL/s with a Nanoject II injector (Drummond scientific) coupled to a stereotaxic frame (Kopf model 902). Optic fibre cannulae (New Doon, FOC-C-1.25-200-7.0-0.37) and stainless steel headplates were affixed to the skull using a combination of light-cured (RelyX Unicem 2, 3M) and self-curing (Superbond, C&B) dental cement.

#### Histological processing

At the end of all behavioural experiments, mice were anaesthetized with intraperitoneally injected pentobarbital (Euthatal or Dolethal 200 mg/mL, 6 mg per 20 g mouse) and transcardially perfused with 10-15 mL of 0.01 M phosphate buffered saline (PBS) followed by 4% paraformaldehyde solution in PBS. Brains were post-fixed at room temperature overnight and cut into 50 *µ*m coronal sections on a vibratome (Leica VT1000 S).

To locate the boundary between the target region (CnF) and its neighbouring pedunculopontine nucleus (PPN), brain slices were immunohistochemically stained for choline acetyltransferase (ChAT). Sections were first permeabilised and blocked in a solution containing 1% bovine serum albumin (BSA, Cambridge Bioscience) and 0.3% Triton X (VWR International Ltd) in 0.01 M PBS for 1.5 hours at room temperature, on a shaker. In parallel, goat anti-ChAT (Millipore AB144P, 0.1 mg/mL, 1:100 dilution) was pre-incubated with an anti-goat Fab fragment (Jackson Immunoresearch AF594, 1.6 mg/mL) in a solution containing 1% BSA and 0.1% Triton in 0.01 M PBS for 1.5 hours at 37^*°*^C The sections were subsequently incubated in this primary antibody solution for 3 days at 37^*°*^C on a shaker. This was followed by three 30-minute washes with the 1% BSA, 0.1% Triton PBS solution and a final 30-minute wash with PBS. Finally, the sections were counterstained and mounted with DAPI Fluoromount-G (Southern Biotech), and imaged at 1.3 *µ*m/px resolution on an epifluorescent microscope (Zeiss Axio Imager 2).

#### Passive treadmill locomotion

After recovery from the surgery, mice were habituated to head fixation for 10 days, 15-50 minutes each day. While head-fixed, mice were allowed to locomote on a low-friction non-motorised treadmill, custom-made based on the documentation and source code released by the Janelia Research Campus [40]. Following the habituation period, 5 second optical stimuli (10-50 Hz, 10 mW at fibre tip) were delivered with a 473 nm laser (Shanghai Laser & Optics Century) for 20-25 trials daily to induce locomotion. Only the mice that showed reliable locomotor responses to the stimulation were used in the experiments. Videos of the left and right side views were recorded at 400 frames per second using a pair of USB3.0 cameras (Basler acA1920-150um). Data acquisition, camera trigger, and optogenetic stimulation were synchronised through a PCIe-6351 board (National Instruments) and controlled using Bonsai [37].

Somatosensory feedback to the animal was manipulated with two types of trials. In head height trials, the height of the head-fixation apparatus was controlled over a 20-25 mm range with ±0.1 mm precision and without any change along the head rotation axes, while keeping the grade fixed at 0 degrees In incline trials, the grade of the treadmill, the head-fixation apparatus, and the cameras was jointly changed in 5±1 deg increments over a [-40, 40] degree range, while keeping the head fixed at a height roughly in the middle of the head height manipulation range. This approximated the “medium” head height used in the incline force sensor experiments.

#### Force sensor measurements

Headplated Vglut2-Cre mice were head-fixed on a set of four aluminium single point load cells (Tedea Huntleigh) such that each cell supported exactly one foot. Head height was modified over a 25-30 mm range with ±0.1 mm precision similar to before. In a separate experiment, setup grade was changed between -40 and 40 degrees at two different head heights. Force measurements were amplified 250000x and low-pass filtered at 100 Hz (pre-amplifier MA103S and amplifier MA102S, both custom-built in the Neuroscience Electronics Lab, University of Cologne), digitised at 1428 Hz (Cambridge Electronic Design Power1401), and recorded in Spike2 software (Cambridge Electronic Design). In parallel, videos with the right side view of the mouse were acquired at 100 Hz using Bonsai [37]. Only the trials with mice standing still for at least 5 seconds were included in analysis.

#### Motorised treadmill locomotion

6-7 weeks old mice were habituated to, and trained on, a motorised treadmill (model 009, custom-built in the Neuroscience Electronics Lab, University of Cologne) for 10 consecutive days. During this period, mice experienced 8-12 locomotor trials per day, each lasting 15 seconds, with the maximum treadmill speed adjusted in a 15-150 cm/s range on an individual, performance-dependent basis using Spike 2 software (Cambridge Electronics Design). Treadmill speed output was used to trigger video data acquisition in a closed-loop manner through a PCIe-6351 board (National Instruments) and Bonsai [37]. Videos from both right and left sides were recorded at 400 frames per second using two USB3.0 cameras (Basler acA1920-150um).

### Quantification and statistical analysis

#### Tracking of body parts

Tracking of the four limbs, snout, back arch, and tail base was performed using DeepLabCut (version 2.1.10.4 [38, 39]). We labelled 20 frames from 72 videos (passive treadmill, each side-view camera separately), 48 videos (force sensor head height trials), 27 videos (force sensor incline trials), or 50 videos (motorised treadmill, each side-view camera separately), and used 95% of those for training. Neural networks were trained for 650000 iterations using ResNet-101 as a starting point, and validated with 10 shuffles. The train and test errors were 3 ± 0 px and 17 ± 2 px respectively (passive treadmill, left side view), 2 ± 0 px and 14 ± 1 px (passive treadmill, right side view), 4.2 ± 0.4 px and 6.1 ± 1.2 px (force sensors, head height trials), 2.2 ± 0.2 px and 5.0 ± 0.5 px (force sensors, incline trials), 2 ± 0 px and 5 ± 0 px (motorised treadmill, left side view), 2 ± 0 px and 7 ± 0 px (motorised treadmill, right side view). For reference, frame size was 1184 × 400 px for both treadmills and 1184 × 900 px for the force sensor setup.

Included in further analysis were only those treadmill trials where at least 90% of the predicted ‘snout’ and ‘hump’ labels and at least 80% of the predicted ‘left hindlimb’ (primary reference limb) labels had a likelihood equal to, or greater than, 0.95.

#### Weight distribution quantification

The fraction of body weight placed on the forelimbs and hindlimbs during standstill trials on the load cells was computed as the ratio between the respective load cell measurements and the total body weight of the mouse. The latter was expressed in terms of voltage through load cell-specific weight-voltage calibration. The difference between the body weight and the cumulative weight supported by all four limbs was attributed to the tensile or reactive force arising from the animal’s interaction with the head-fixation apparatus. Body weights were empirically determined within 0-3 days of load cell data acquisition and interpolated in-between those measurements.

In head height trials, the absolute head heights were rescaled and divided by weight such that a weight-adjusted head height value of 1 reflected the maximum height at which the forefeet were in full contact with the ground.

The two-dimensional centre of gravity was computed from the normalised load cell data as follows:

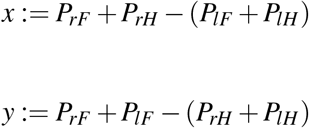

where rF, rH, lF, and lH represent measurements from the load cells that support the respective limbs.

#### Gait characterisation

Gait was characterised by estimating the phases of limb movement relative to a reference limb, typically the left or right hindlimb. Analysis was focused on the time periods where the speed of the self-paced or motorised treadmill belt exceeded 1 cm/s. Stance and swing onsets for the reference limb were identified by applying a peak detection algorithm to the respective x coordinate time series, with consecutive stance-to-stance intervals being treated as strides. For every stride of duration *d*, the x coordinate time series of each non-reference limb was cross-correlated with that of the reference limb over for delays in range 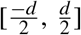 to reveal an interlimb phase difference. This was subsequently normalised by stride duration yielding a value in interval [−0.5, 0.5] or [*−π, π*] in polar coordinates. If the phase values of two independently tracked points on the same foot differed by more than 0.05, the particular stride was excluded from analysis.

Given the lack of clear boundaries between the observed combinations of limb phases [41], the analysis refrained from assigning discrete gait labels to individual strides and instead considered limb phase distributions under various conditions. Density distributions of limb phases excluded data from mice that performed fewer than 20 strides in a given category. Other analyses did not exclude data on the basis of sparseness.

Pairwise similarity between the discrete density distributions across categories, as well as their similarity to uniform distributions and ones corresponding to idealised trot and bound, was quantified through the circular optimal transport (COT) [42]. In short, given two probability distributions *µ* and *v*, COT reflects the minimum cost of transporting the mass from one distribution to the other by taking into account the geometry of the space, as:

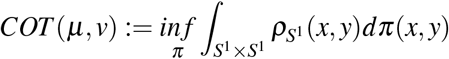

where *S*^1^ is a circle parametrised by [0, 1) and equipped with geodesic distance

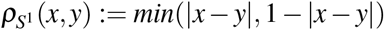

Since COT is a symmetrical measure, the upper-diagonal values of the pairwise COT matrices were not plotted. Statistical significance of COTs was assessed using a permutation test with 10000 repetitions and the correspondence between limb phase and snout-hump angle or treadmill grade – randomly shuffled within an animal. Bonferroni correction was applied to account for multiple comparisons.

#### Mixed-effects models

To quantify the contributions of various fixed predictors, including locomotor speed, body tilt angle, treadmill incline, age, weight, and sex, to the relative limb phase series of Bayesian mixed-effects regression models for circular data were fitted using the *bpnme* function from the *bpnreg* package (version 2.0.2, [20]). Mouse identity was incorporated into the models as a random intercept. The models were fitted using random 50% of the data due to a large dataset size, with 100 burn-in iterations, a total of 1000 output iterations, and a lag of 3. Model convergence was verified with traceplots for all predictors (not shown), and the significance of individual continuous predictors was assessed using the highest posterior density (HPD) interval of the signed shortest distance to origin (SSDO) as described in [20]. Statistical significance between levels of categorical variables was also evaluated based on the respective HPD intervals. Model comparison was based on the Watanabe-Akaike information criterion (WAIC).

Relationships between non-circular variables were quantified with linear mixed-effects models using the *lmer* function from the *lme*4 package (version 1.1-28, [43]). As in the circular models, mouse identity was modelled as a random intercept. Predictors with variance inflation factor greater than 5 were excluded from the models due to collinearity, although centering of the variables often resolved this problem. The significance of individual predictors was evaluated based on t-tests with Satterthwhite approximation, while model comparison relied on the Akaike information criterion (AIC).

All models referenced in this paper that feature a linear outcome variable are shown in Table 1, and those with a circular outcome variable are shown in Table 2. Simulation of idealised canonical gaits To ground the observed limb phase data in the canonical gait classification framework [3, 5, 6], combinations of limb phases representing idealised trot, bound, as well as transverse and rotary gallop were simulated as a set of normal distributions with a standard deviation of 0.06*π*. The phase distributions of trot were centred at (0.5*π*, 0.5*π*, 0) for the homologous, homolateral, and diagonal limb respectively. Analogously, the phase distributions for bound, transverse gallop, and rotary gallop were centred at (0, −0.4*π, −*0.4*π*), (0.15*π*, 0.3*π, −*0.3*π*), and (0.15*π, −*0.3*π*, 0.3*π*).

#### Computation of GRF-equivalent shifts in limb phase

To compare the phase shifts observed in the two types of trials, it was assumed that a given change in treadmill incline on animal head height resulted in the same average weight redistribution on the non-motorised treadmill as it did on the force sensors (medium head height in the incline trials) where it was directly measured. The validity of this assumption is discussed in the paper.

With that in mind, we first determined the population-averaged anteroposterior shift in the relative GRF distribution that corresponds to the range of snout-hump angles seen in the incline trials. This was followed by finding the incline range corresponding to the previously identified shift in the anteroposterior GRF distribution and using that to determine the GRF-equivalent limb phase shift due to a change in treadmill incline. The same range of snout-hump angles was then used to find the GRF-equivalent shift in limb phase in the head height trials.

These GRF-equivalent phase shifts were computed on a per-mouse basis using (1) individual animal data from the treadmill experiments because the same set of mice performed both head height and incline trials; (2) population-averaged force sensor data because few of the same mice were used in the forces sensor experiments.

#### Histological quantification

Brain sections were registered to the Allen Mouse Brain Common Coordinate Framework (CCFv3, [44]) using the ABBA plugin in ImageJ (BioImaging And Optics Platform, Ecole Polytechnique Federale de Lausanne). The positions of virally labelled cells and optic fibre implants were subsequently determined in Qupath [45] using its cell detection and manual annotation tools respectively. Finally, the data were visualised in 3D with Brainrender [46].

## Supplementary information

**Figure S1.**
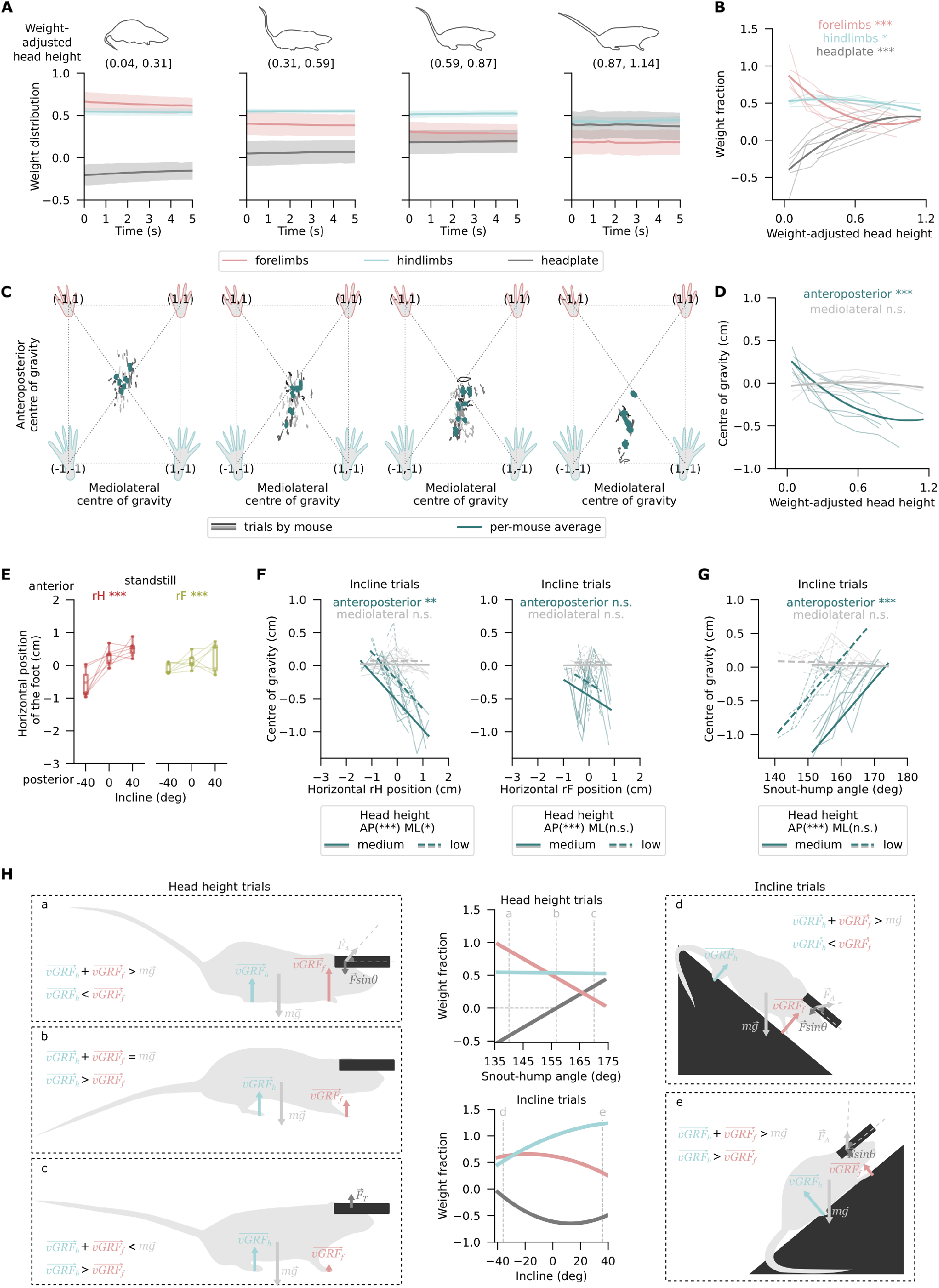
Head height modulates the relative fore/hind GRF in a curvilinear manner, and mice oppose incline-related GRF redistribution through postural changes. **(A)** The average fraction of body weight placed on forelimbs, hindlimbs, and the head fixation apparatus (pink, blue, and grey solid lines respectively) over 5 second standstills across mice (n=7), grouped by weight-adjusted head height. The shaded areas show 95% confidence intervals. **(B)** Same data as in (A), but shown as trial averages of individual mice computed over four discrete ranges of weight-adjusted head heights (thin lines), along with second-order polynomial mixed-effects model fits (thick; statistics in Table S1, row 1). **(C)** The centre of gravity across limbs during the same head-height trials as in (A), grouped by weight-adjusted head height. Grey traces show individual trials with shade indicative of mouse identity, while per-mouse means are displayed in teal. **(D)** Same as (C) but shown as trial averages of individual mice (thin lines) and second-order polynomial mixed-effects model fits (thick; statistics in Table S1, row 1) for the mediolateral (grey) and anteroposterior (teal) body axes separately. **(E)** Anteroposterior position of the right hindlimb and forelimb during standstill trials on load cells at three inclines, averaging across the two head heights. Per-mouse means across inclines are shown as connected dots, whereas boxplots summarise data across mice. **(F)** The centre of GRF distribution along the mediolateral (grey) and anteroposterior (teal) body axes during incline trials at a higher (solid) and lower (dashed) head height, shown as a function of anteroposterior position of the right hindlimb (left) and right forelimb (right). Data from individual mice are plotted as thin lines, whereas the thick lines show linear mixed effects model fits (statistics in Table S1, rows 3 and 4). **(G)** Same as (F) but plotted as a function of the snout-hump angle (statistics in Table S1, rows 5). **(H)** Illustration of the vertical GRFs and other vertical forces (not measured) acting on, or generated by, the animal at different head heights (a, b, c) and inclines (d, e). Statistical significance thresholds: * *p <* 0.05, ** *p <* 0.01, *** *p <* 0.001.

**Figure S2.**
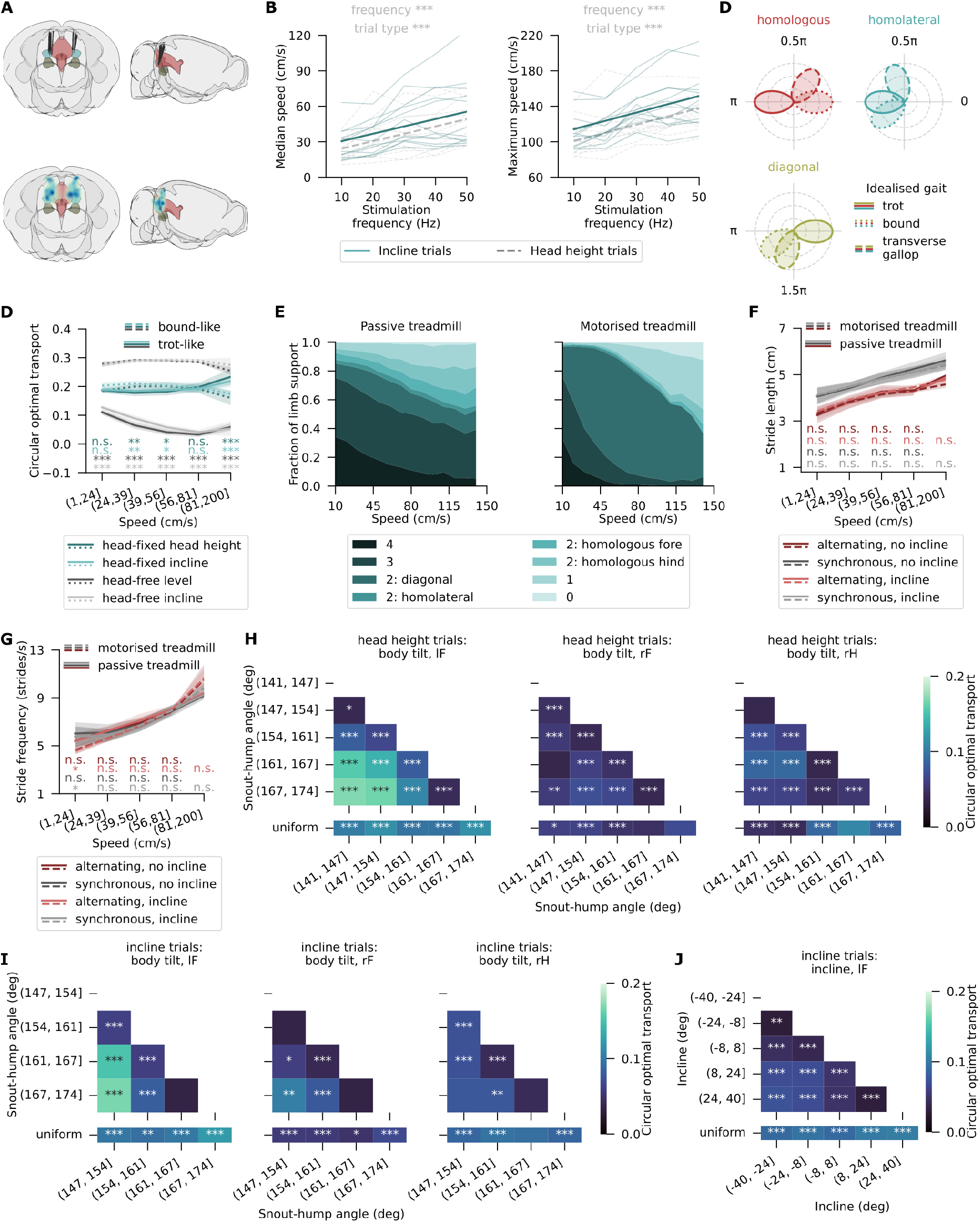
Optogenetically evoked head-fixed locomotion resembles natural head-free locomotion and reveals sensory feedback-sdependent effects on homolateral coordination. **(A)** Traced positions of optic fibre implants (*top*) and density distributions of virally labelled cells (*bottom*) viewed in the coronal (*left*) and sagittal (*right*) plane. Cuneiform nucleus, pedunculopontine nucleus, and periaqueductal grey are shown in teal, dark yellow, and pink respectively. **(B)** Median speed (*left*, slope: 0.62± 0.04 cm/s per Hz, incline effect: 6.3± 1.3 cm/s) and maximum speed (*right*, slope: 0.94± 0.07 cm/s per Hz, incline effect: 13.7± 2.2 cm/s) of optogenetically evoked head-fixed locomotion depending on the optical stimulation frequency, shown for individual mice (thin lines) and as linear mixed-effects regression across mice (thick lines). **(C)** Simulated phases of non-reference limbs, characteristic of an idealised trot (solid lines), bound (dotted), and transverse gallop (dashed) with the homologous as the leading leg. **(D)** Circular optimal transport costs [42] to transform the observed limb phase distributions into the simulated distributions of idealised trot and bound from (D) in speed quintiles, with the costs averaged across limbs and mice. As in (D), data from optogenetically induced head-fixed locomotion at variable incline and head height is shown in light and dark teal respectively, whereas the grey traces represent head-free locomotor data acquired on a motorised treadmill at zero (dark grey) or variable (light grey) incline. Shaded regions are 95% confidence intervals around the mean value across mice. **(E)** Area plot of average foot support types as a fraction of stride cycle across speeds for passive treadmill (*left*) and motorised treadmill (*right*), based on one thousand randomly selected strides per mouse and inspired by [47]. **(F)** Length of strictly alternating (light/dark red) and strictly synchronous (light/dark grey) strides in speed quintiles, averaged across mice. Solid lines represent optogenetically induced head-fixed locomotion in head height (dark red/grey) and incline (light red/grey) trials, whereas dashed lines are from head-free motorised treadmill locomotion at zero (dark red/grey) and variable (light red/grey) incline. **(G)** Same as (F) but showing stride frequency. **(H)** Circular optimal transport costs to transform lF, rF, and rH phase distributions relative to lH across snout-hump angles (data from head height trials, Figure 2D) into one another (*top row*), or into a uniform distribution (*bottom row*), with Bonferroni-corrected two-tailed permutation tests. **(I)** Same as (H) but with data from incline trials (also in Figure 2D). **(J)** Same as (H) but only for lF and across treadmill inclines (data from Figure 3A). Statistical significance thresholds: * *p <* 0.05, ** *p <* 0.01, *** *p <* 0.001.

**Figure S3.**
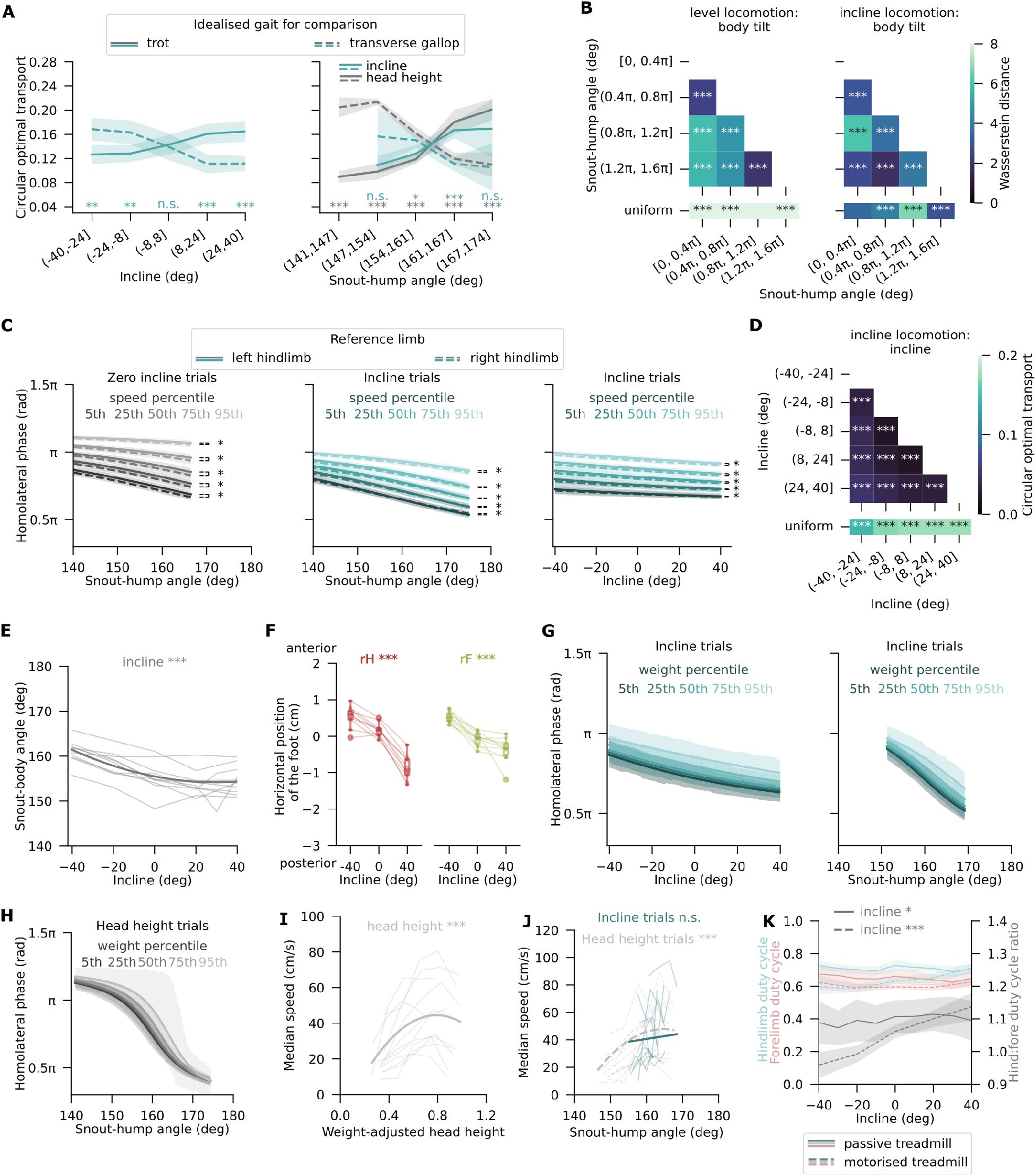
Head-fixed and head-free locomotion display similar sensory feedback-dependence of homolateral coordination. **(A)** Circular optimal transport cost to transform the homolateral limb phase distributions from Figure 2D (*left*) and Figure 3A (*right*) into simulated phase distributions corresponding to idealised trot (solid lines) and transverse gallop (dashed lines). Shaded regions are 95% confidence intervals. **(B)** Circular optimal transport costs to transform the relative lF phase distributions across snout-hump angles during motorised treadmill locomotion (data from Figure 4B) into one another (*top row*), or into a uniform distribution (*bottom row*), with Bonferroni-corrected two-tailed permutation tests. **(C)** Forelimb phases relative to the homolateral hindlimb as a function of motorised treadmill incline or snout-hump angle with lH (solid lines) and rH (dashed lines) as the reference limb. Shown are Bayesian mixed-effects model fits to the incline trial data at five percentiles of speed. Shaded regions are 95% highest posterior density intervals. **(D)** Same as (B) but for head-free locomotion on zero (*left*) or variable (*middle, right*) inclines. **(E)** Body tilt angle as a function of treadmill incline during zero incline, plotted for individual mice across trials (thin lines) and as linear mixed-effects model fits across mice (thick). **(F)** Horizontal foot position depending on the treadmill incline during motorised treadmill locomotion. **(G)** Same as (C) but with passive treadmill data from incline trials at five percentiles of animal body weight. **(H)** Same as (G) but showing data from the head height trials. **(I)** Median speed of passive treadmill locomotion plotted against weight-adjusted head height for individual mice (thin lines) and as linear mixed-effects regression across mice (thick lines). **(J)** Same as (I) but as a function of snout-hump angle, showing data from incline (teal, solid) and head height (grey, dashed) trials. **(K)** Average hindlimb (teal) and forelimb (pink) duty cycles, and their ratio (grey) during head-fixed (*p*_*ratio*_ = 0.04, one sample t-test) and head-free (*p*_*ratio*_ = 5.9 × 10^−15^) locomotion, across treadmill inclines. Mean values across mice are shown, along with 95% confidence intervals. Statistical significance in Bayesian mixed-effects models (C,G,H): * significant, n.s. not significant. Statistical significance thresholds: * *p <* 0.05, ** *p <* 0.01, *** *p <* 0.001.

Table S1: Linear and circular mixed-effects models fit to the data and referenced in the paper. 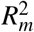 and 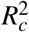 are the marginal and conditional *R*^2^ estimates, reflecting the variance explained by the fixed effects alone or through both fixed and random effects.

